# RIPK3 promotes brain region-specific interferon signaling and restriction of tick-borne flavivirus infection

**DOI:** 10.1101/2023.01.23.525284

**Authors:** Marissa Lindman, Juan P Angel, Irving Estevez, Nydia P Chang, Tsui-Wen Chou, Micheal McCourt, Colm Atkins, Brian P. Daniels

## Abstract

Innate immune signaling in the central nervous system (CNS) exhibits many remarkable specializations that vary across cell types and CNS regions. In the setting of neuroinvasive flavivirus infection, neurons employ the immunologic kinase receptor-interacting kinase 3 (RIPK3) to promote an antiviral transcriptional program, independently of the traditional function of this enzyme in promoting necroptotic cell death. However, while recent work has established roles for neuronal RIPK3 signaling in controlling mosquito-borne flavivirus infections, including West Nile virus and Zika virus, functions for RIPK3 signaling in the CNS during tick-borne flavivirus infection have not yet been explored. Here, we use a model of Langat virus (LGTV) encephalitis to show that RIPK3 signaling is specifically required in neurons of the cerebellum to control LGTV replication and restrict disease pathogenesis. This effect did not require the necroptotic executioner molecule mixed lineage kinase domain like protein (MLKL), a finding similar to previous observations in models of mosquito-borne flavivirus infection. However, control of LGTV infection required a unique, region-specific dependence on RIPK3 to promote expression of key antiviral interferon-stimulated genes (ISG) in the cerebellum. This RIPK3- mediated potentiation of ISG expression was associated with robust cell-intrinsic restriction of LGTV replication in cerebellar granule cell neurons. These findings further illuminate the complex roles of RIPK3 signaling in the coordination of neuroimmune responses to viral infection, as well as provide new insight into the mechanisms of region-specific innate immune signaling in the CNS.

**Importance:** Interactions between the nervous and immune systems are very carefully orchestrated in order to protect the brain and spinal cord from immune-mediated damage, while still maintaining protective defenses against infection. These specialized neuro-immune interactions have been shown to vary significantly across regions of the brain, with innate antiviral signaling being particularly strong in the cerebellum, although the reasons for this are poorly understood. Here, we show a specialized adaptation of programmed cell death signaling that uniquely protects the cerebellum from tick-borne flavivirus infection. These findings provide important new insight into the molecular mechanisms that promote the uniquely robust antiviral immunity of the cerebellum. They also provide new clues into the pathogenesis of tick-borne encephalitis, a zoonosis of significant global concern.

## Introduction

Flaviviruses are a family of positive sense RNA viruses which include several notable pathogens associated with neuroinvasive infection in humans, including West Nile virus (WNV), Zika virus (ZIKV), and Japanese Encephalitis virus (1). While nearly all major flaviviruses are transmitted by mosquito vectors, a small but significant number of flaviviruses are transmitted by ticks, including Tick-borne encephalitis virus (TBEV) and its close relatives that together make up a single TBEV serocomplex. Tick borne encephalitis is a significant and growing threat to public health, particularly in Europe and northern Asia, where TBEV constitutes the most prevalent tick-borne zoonotic disease (2–4). Notably, some TBEV strains elicit mortality rates up to 40% in humans (5), underscoring the urgent need to better understand the mechanisms underlying the pathogenesis of tick-borne flavivirus infections.

Effective control of flavivirus infection in the central nervous system (CNS) requires robust innate immune signaling in neural cells, particularly neurons, which are the predominantly infected cell type in most cases of flavivirus encephalitis (6–9). Effective type I interferon (IFN) signaling is of particular importance for innate control of viral replication in neurons (10–12). Notably, differences in type I IFN signaling across neural cell types and brain regions are associated with differential susceptibility to flavivirus infection. For example, previous reports suggest that the enhanced type I IFN signaling observed in hindbrain regions compared to the forebrain is an underlying determinant of the enhanced susceptibility of forebrain regions to WNV infection (12, 13). However, the unique signaling mechanisms that promote differential IFN-mediated control of viral infection in the hindbrain have not been extensively characterized.

A potential regulator of neuronal IFN signaling during flavivirus infection is receptor interacting protein kinase-3 (RIPK3). RIPK3 is an enzyme traditionally associated with necroptosis, a form of lytic programmed cell death (14). Necroptosis occurs via the RIPK3- dependent activation of mixed lineage kinase domain like protein (MLKL), which forms oligomeric pore complexes that induce cellular lysis (15). However, many recent studies have identified complex roles for RIPK3 signaling in the coordination of inflammation, including the regulation of inflammatory transcriptional responses that occur independently of necroptosis (16–24). We and others have demonstrated that RIPK3 signaling in neurons is of particular importance for the control of neurotropic viral infections, as neuronal RIPK3 promotes a robust antimicrobial transcriptional program, including many IFN stimulated genes (ISGs), that restricts viral infection without inducing neuronal necroptosis (16, 17). Other recent studies have identified unexpected roles for RIPK3 in the regulation of type I IFN signaling, via mechanisms which include the regulation of pattern recognition receptor signaling and protein kinase-R (PKR)-mediated stabilization of *Ifnb* mRNA (18, 19).

In this study, we interrogated roles for RIPK3 in controlling tick-borne flavivirus infection. To do so, we used Langat virus (LGTV), a naturally attenuated member of the TBEV serocomplex that can be studied under BSL2 containment. *Ripk3*^-/-^ mice exhibited enhanced clinical disease following subcutaneous LGTV infection, while *Mlkl*^-/-^ mice were indistinguishable from littermate controls, suggesting a necroptosis-independent function for RIPK3 in restricting LGTV pathogenesis. Notably, *Ripk3*^-/-^ mice exhibited increased viral burden in the cerebellum, along with diminished expression of inflammatory chemokines and ISGs in the cerebellum, but not the cerebral cortex. *In vitro* analysis of cultured primary cortical and cerebellar cell types showed that pharmacologic inhibition of RIPK3 resulted in enhanced LGTV replication in cerebellar granule cell neurons but not in cortical neurons or in astrocytes derived from either brain region. Transcriptional profiling showed that RIPK3 signaling was uniquely required for the full induction of ISG expression in cerebellar granule cell neurons, demonstrating a previously unknown, region-specific function for RIPK3 in coordinating innate antiviral immunity within the CNS.

## Results

### RIPK3 controls LGTV pathogenesis independently of MLKL and peripheral immunity

To assess the role of RIPK3 in controlling LGTV pathogenesis, we subcutaneously infected *Ripk3*^-/-^ mice, along with heterozygous littermate controls, with 3×10^4^ plaque forming units (pfu) of the Malaysian LGTV strain TP21. We note that *Ripk3*^+/-^ animals do not exhibit haploinsufficiency and are routinely used as littermate controls in studies of this pathway (25–27). Control animals exhibited limited mortality following LGTV infection (Figure 1A), consistent with previous reports (28, 29). However, mice lacking *Ripk3* expression exhibited a significantly accelerated and enhanced rate of mortality (Figure 1A). In addition, a higher proportion of *Ripk3*^-/-^ mice exhibited clinical signs of neurologic disease, including paresis or full hindlimb paralysis, by 14 days post infection (dpi) (Figure 1B), and this difference persisted to at least 21 dpi. These data suggest that *Ripk3* is essential for restricting neuropathogenesis during LGTV infection.

**Figure 1.**
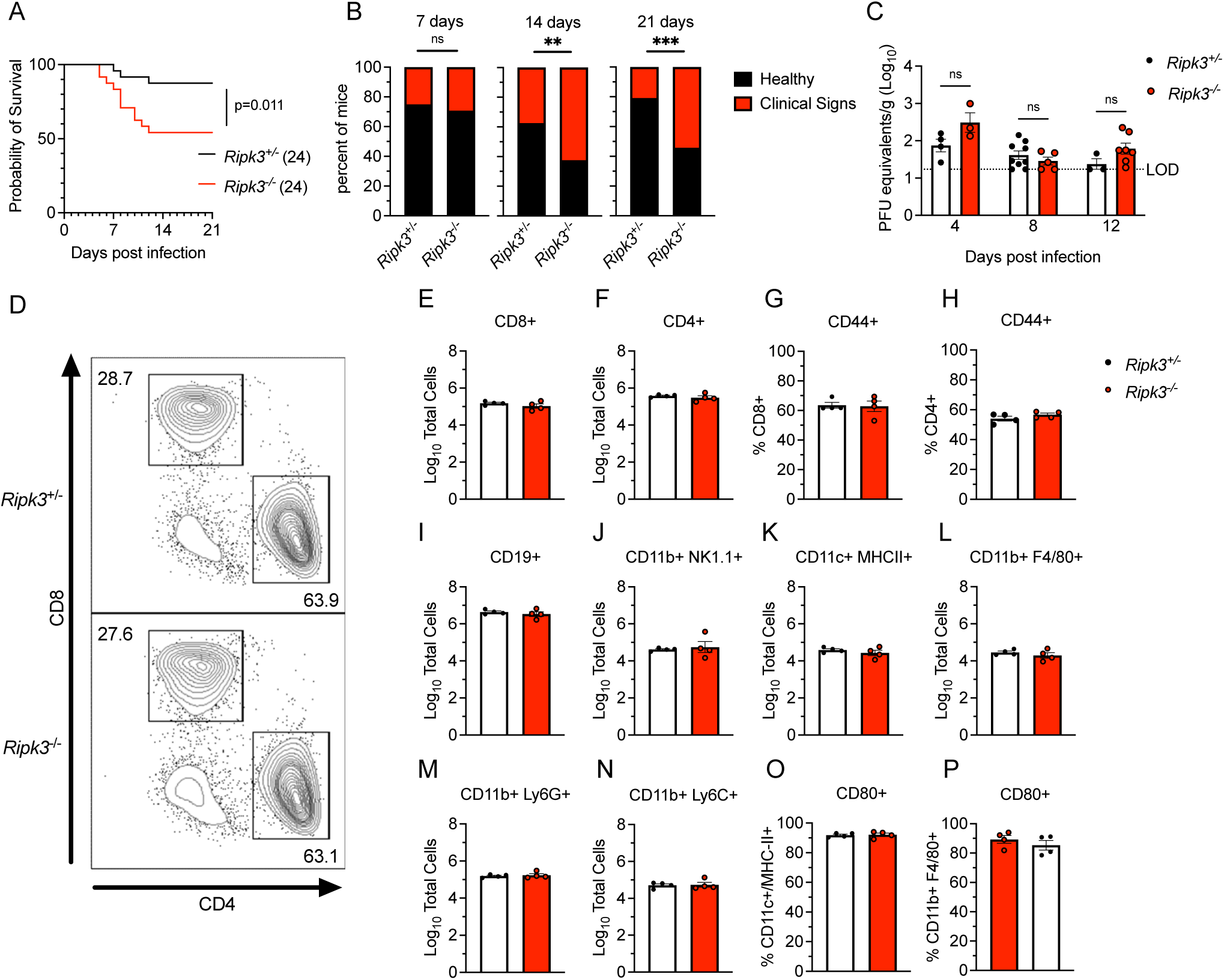
RIPK3 limits LGTV pathogenesis independently of peripheral immunity. (A-B) Survival analysis (A) and presentation of clinical signs of disease (B) in *Ripk3*^-/-^ mice and littermate controls following subcutaneous inoculation with 3×10^4^ PFU LGTV TP21. Data are pooled from two experiments. (C) *Ripk3*^-/-^ and littermate control mice were infected subcutaneously with LGTV TP21. On indicated days following infection, splenic viral burden was measured via qRT-PCR. Data was normalized against a standard curve of known viral titers to generate plaque-forming unit (PFU) equivalents. Data for each day post infection are pooled from 2-3 experiments. LOD, limit of detection. (D-P) *Ripk3*^-/-^ and littermate control mice were infected subcutaneously with LGTV TP21 for 8 days prior to harvesting splenocytes and profiling leukocytes by flow cytometry. (D) Representative flow cytometry plots showing CD8+ and CD4+ T cells among CD3+ leukocytes in the spleen. Numbers represent percentage of cells in each gate relative to total plotted cells. (E-F) Numbers of CD8+ T cells (E) and CD4+ T cells (F) among CD3+ leukocytes. (G-H) Percentage of CD44+ cells among CD8+ T cells (G) and CD4+ T cells (H). (I-N) Numbers of CD19+ B cells (I), CD11b+ NK1.1+ Natural Killer cells (J), CD11c+ MHC-II+ dendritic cells (K), CD45high CD11b+ F4/80+ macrophages (L), CD11b+ Ly6G+ neutrophils (M), and CD45high CD11b+ Ly6C+ monocytes (N) among total leukocytes in the spleen. (O-P) Percentage of CD80+ cells among CD11c+ MHC-II+ dendritic cells (O) and CD11b+ F4/80+ macrophages (P). ns, not significant. **p < 0.01, ***p < 0.001. Error bars represent SEM.

To better understand this phenotype, we first assessed whether RIPK3 was required for early control of systemic infection. Spleens of infected mice exhibited low levels of LGTV RNA that were not impacted by *Ripk3* expression (Figure 1C). To test whether *Ripk3*^-/-^ mice exhibited any deficiencies in peripheral immune responses, we performed flow cytometric analysis of major immune cell subsets in the spleens of infected animals at 8 dpi. *Ripk3*^-/-^ animals exhibited similar frequencies (Figure 1D) and total numbers (Figure 1E-F) of CD4 and CD8 T cells among all splenocytes compared to littermate controls, as well as similar rates of CD44 expression (a key T cell activation marker) across both subsets (Figure 1G-H). Numbers of B cells (Figure 1I) and natural killer (NK) cells (Figure 1J) were also similar between genotypes. In the myeloid compartment, we observed similar numbers of CD11c^+^ MHCII^+^ dendritic cells (Figure 1K) between genotypes, as well as similar numbers of myeloid subsets expressing F4/80 (Figure 1L), Ly6G (Figure 1M), and Ly6C (Figure 1N). Both CD11c^+^ MHCII^+^ and F4/80^+^ antigen presenting cell subsets also exhibited similar rates of expression of the costimulation signal CD80 between genotypes (Figure 1O-P). These data suggest that *Ripk3*^-/-^ mice mounted normal peripheral immune responses to subcutaneous LGTV challenge, similar to our previous observations with WNV and ZIKV (16, 17). Thus, the increased pathogenesis observed in mice lacking *Ripk3* was unlikely to arise from a failure in peripheral virologic control.

A potential mechanism by which RIPK3 signaling might restrict LGTV pathogenesis is through the induction of necroptosis in infected cells. We thus tested whether loss of the necroptotic executioner molecule MLKL would impact disease course following subcutaneous LGTV infection. Notably, *Mlkl*^-/-^ mice exhibited no difference in either survival or development of clinical disease signs compared to littermate controls (Figure 2A-B). We saw similarly that *Mlkl*^-/-^ did not exhibit altered splenic viral burden at 8dpi (Figure 2C). Flow cytometric analysis also revealed essentially identical numbers and frequencies of all major immune cell subsets in the spleen at this time point (Figure 2C-P). Multistep growth curve analysis also demonstrated that neither RIPK3 nor MLKL impacted the low levels of LGTV replication observed in primary leukocyte cultures, including bone marrow derived macrophages and dendritic cells (Supplemental Figure 1A-B). These data suggest that MLKL, and therefore necroptosis, is not a major contributor to peripheral virologic control or overall disease pathogenesis in the setting of LGTV infection, and thus that RIPK3 exerts its protective effect in this model through an alternative mechanism.

**Figure 2.**
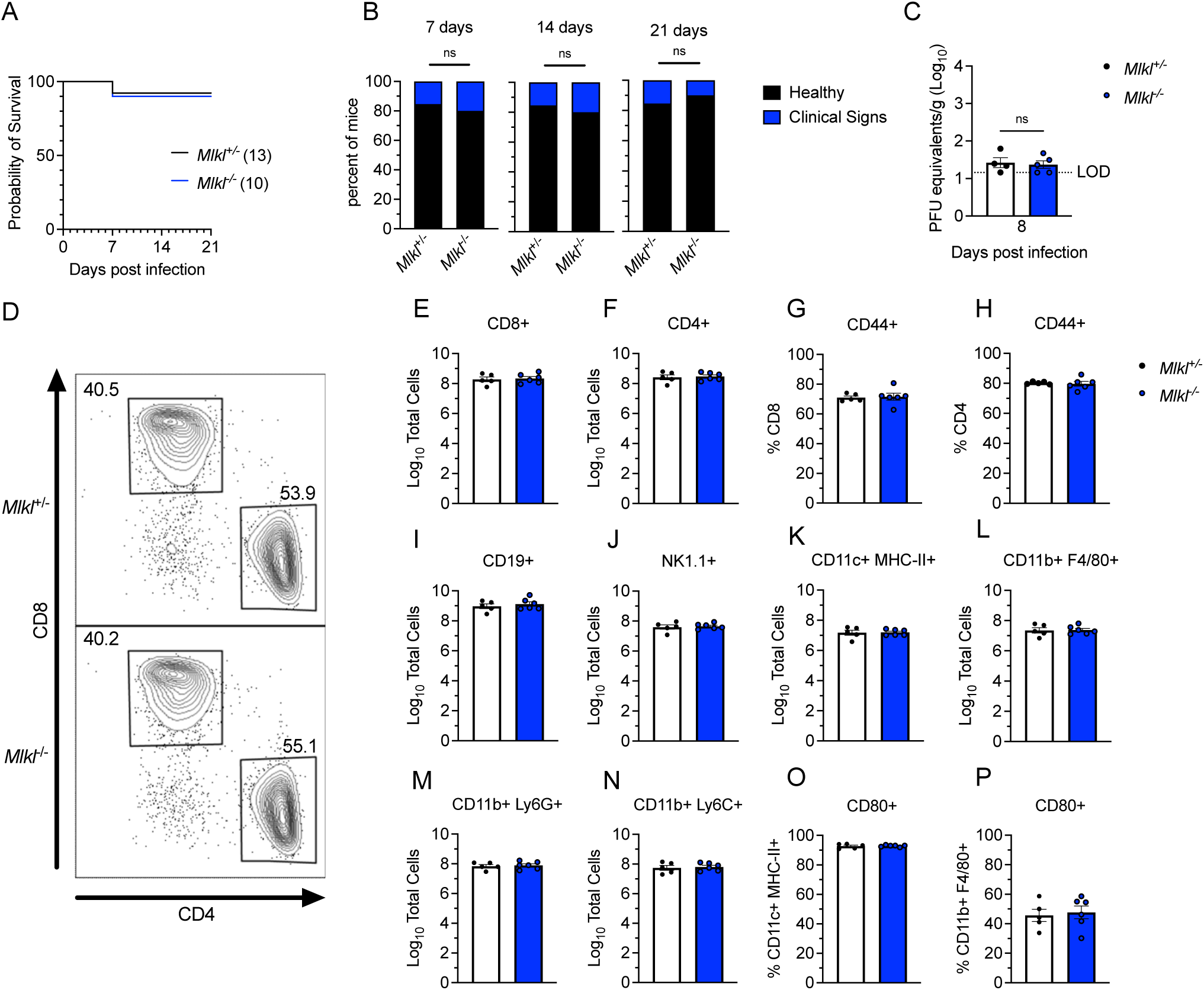
MLKL signaling does not influence Langat virus pathogenesis. (A-B) Survival analysis (A) and presentation of clinical signs of disease (B) in *Mlkl*^-/-^ mice and littermate controls following subcutaneous inoculation with 3×10^4^ PFU LGTV TP21. Data are pooled from two experiments. (C) *Mlkl*^-/-^ and littermate control mice were infected subcutaneously with LGTV TP21. On indicated days following infection, splenic viral burden was measured via qRT-PCR. Data was normalized against a standard curve of known viral titers to generate plaque-forming unit (PFU) equivalents. Data for each day post infection are pooled from 2-3 experiments. LOD, limit of detection. (D-P) *Mlkl*^-/-^ and littermate control mice were infected subcutaneously with LGTV TP21 for 8 days prior to harvesting splenocytes and profiling leukocytes by flow cytometry. (D) Representative flow cytometry plots showing CD8+ and CD4+ T cells among CD3+ leukocytes in the spleen. Numbers represent percentage of cells in each gate relative to total plotted cells. (E-F) Numbers of CD8+ T cells (E) and CD4+ T cells (F) among CD3+ leukocytes. (G-H) Percentage of CD44+ cells among CD8+ T cells (G) and CD4+ T cells (H). (I-N) Numbers of CD19+ B cells (I), CD11b+ NK1.1+ Natural Killer cells (J), CD11c+ MHC-II+ dendritic cells (K), CD45high CD11b+ F4/80+ macrophages (L), CD11b+ Ly6G+ neutrophils (M), and CD45high CD11b+ Ly6C+ monocytes (N) among total leukocytes in the spleen. (O-P) Percentage of CD80+ cells among CD11c+ MHC-II+ dendritic cells (O) and CD11b+ F4/80+ macrophages (P). ns, not significant. Error bars represent SEM.

### RIPK3 is required for CNS-intrinsic restriction of LGTV infection

Because we did not observe differences in peripheral virologic control in *Ripk3*^-/-^ mice, we next questioned whether RIPK3 acted in a CNS-intrinsic manner to limit LGTV infection. To assess this, we next used an intracranial infection route in order to assess local effects of RIPK3 signaling on LGTV pathogenesis. *Ripk3*^-/-^ mice exhibited accelerated and enhanced mortality compared to littermate controls following intracranial infection (Figure 3A). *Ripk3*-deficient mice also exhibited worsened clinical disease prior to death, as evidenced by earlier and more dramatic weight loss following infection (Figure 3B). In contrast, *Mlkl*^-/-^ mice were indistinguishable from littermate controls in terms of overall mortality (Figure 3C) and weight loss (Figure 3D) following intracranial infection. These data further supported the idea that RIPK3 restricts LGTV neuropathogenesis via CNS-intrinsic mechanisms, independently of necroptosis.

**Figure 3.**
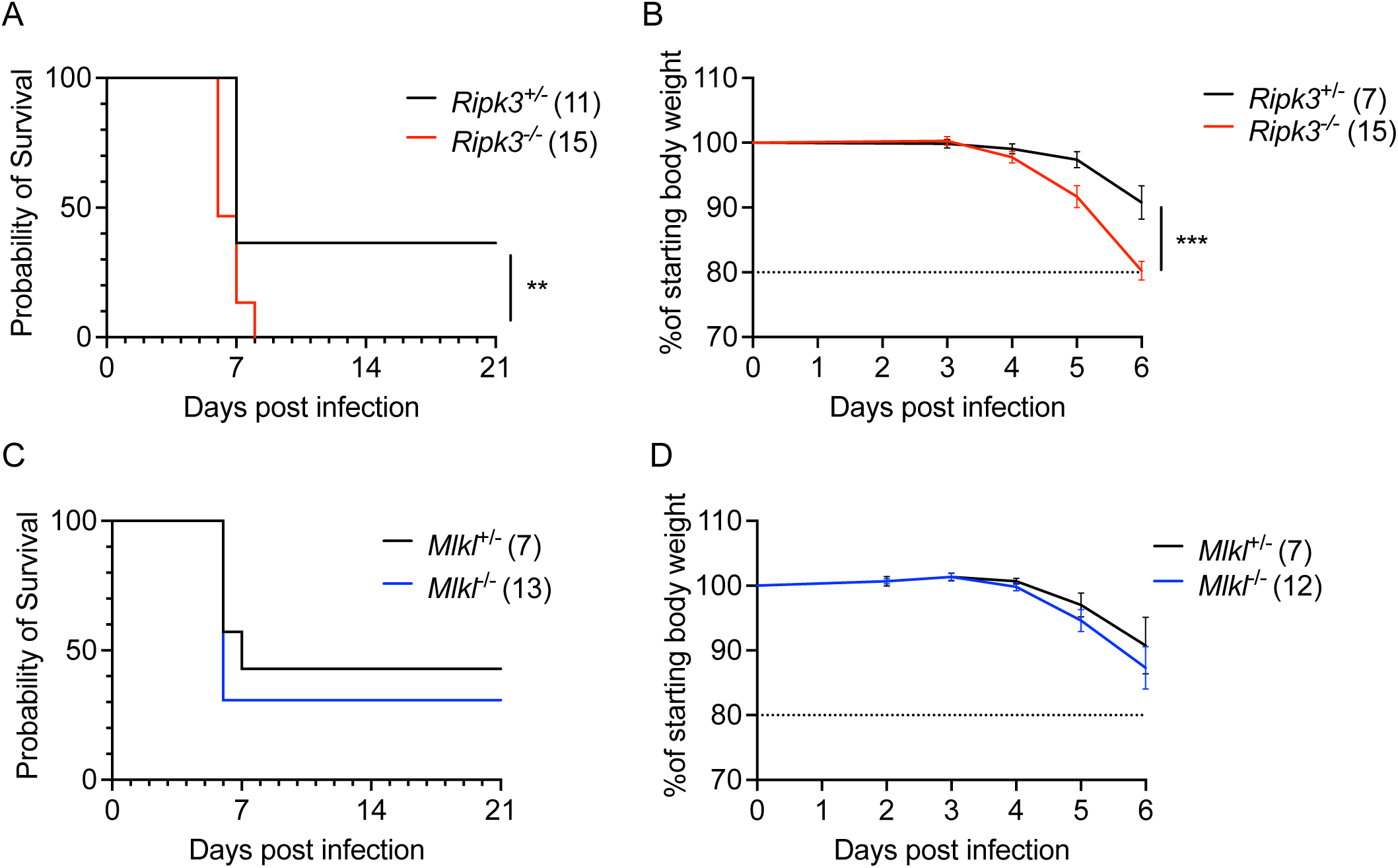
RIPK3, but not MLKL, restricts Langat virus pathogenesis following intracranial infection. Survival and body weight analysis from *Ripk3*^-/-^ (A-B) and *Mlkl*^-/-^ (C-D) mice and their respective littermate controls following intracranial inoculation with 50 PFU LGTV TP21. Data are pooled from two (A-B) or three (C-D) experiments. ns, not significant. **p < 0.01, ***p < 0.001.

### RIPK3 promotes neuronal chemokine expression in a region-specific manner following LGTV infection

We and other previously showed that neuronal RIPK3 signaling was required for the expression of key inflammatory chemokines that served to restrict WNV pathogenesis by coordinating the recruitment of leukocytes into the infected CNS. We thus questioned whether RIPK3 also promotes chemokine expression in the CNS during LGTV infection. Surprisingly, transcriptional profiling in the cerebral cortex of *Ripk3*^-/-^ mice following subcutaneous LGTV infection revealed no differences in expression of major chemokines compared to littermate controls (Figure 4A). However, we did observe significantly diminished chemokine responses in cerebellar tissues derived from *Ripk3*^-/-^ animals (Figure 4B). To understand which cell types were driving this region-specific deficit in chemokine expression, we next cultured primary neurons and astrocytes derived specifically from either cerebral cortex or cerebellum and infected with LGTV, with or without a small molecule inhibitor of RIPK3 (GSK 872). Consistent with our *in vivo* findings, blockade of RIPK3 in cerebral cortical neurons did not impact chemokine expression following LGTV infection (Figure 4C). In contrast, infected cerebellar granule cell neuron cultures exhibited significantly diminished chemokine expression when RIPK3 was inhibited by GSK 872 (Figure 4D). Notably, we did not observe a dependence on RIPK3 for the expression of chemokines in astrocytes derived from either region (Figure 4E-F). These data suggest that RIPK3 serves an unexpected, region-specific transcriptional function in neurons of the cerebellum during neuroinvasive LGTV infection.

**Figure 4.**
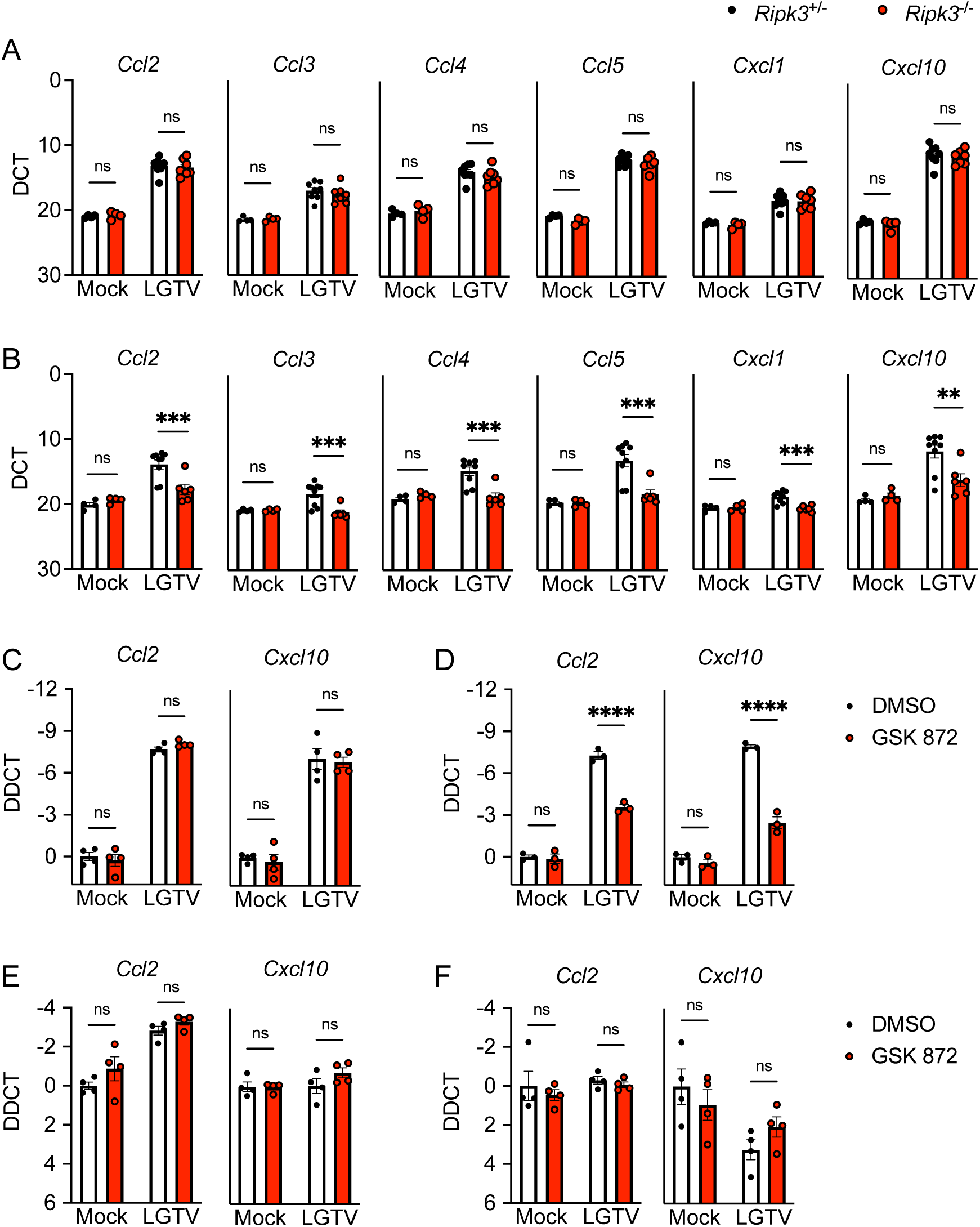
RIPK3 promotes chemokine expression in the cerebellum during LGTV encephalitis. (A-B) *Ripk3* ^-/-^ and littermate control mice were infected subcutaneously with LGTV TP21. At 8dpi cerebral cortical (A) and cerebellar tissues (B) were harvested and assayed for chemokine transcripts via qRT-PCR. (C-F) *Ccl2* and *Cxcl10* expression in wildtype (C57BL/6J) cultures of primary cortical neurons (C), cerebellar granule cell neurons (D), cortical astrocytes (E), and cerebellar astrocytes (F) following 2-hour pretreatment with GSK872 or vehicle and 24h infection with 0.5 (C-D) or 0.01 (E-F) MOI LGTV TP21, measured via qRT-PCR. ns, not significant. *p<0.05, **p < 0.01, ***p < 0.001, ****p < 0.0001. Error bars represent SEM.

### RIPK3 is not required for immune cell recruitment to the LGTV-infected CNS

We next questioned whether diminished chemokine expression in the cerebellum of *Ripk3*^-/-^ mice would result in a failure to recruit antiviral leukocytes into this brain region. We thus performed flow cytometric analysis of leukocytes derived from either cerebral cortex or cerebellum following subcutaneous LGTV infection. Remarkably, we saw no evidence of changes in lymphocyte recruitment in either brain region of *Ripk3*^-/-^ mice compared to littermate controls on either 6 or 8 dpi (Figure 5A). This lack of difference extended across all major CD45^hi^ infiltrating leukocyte subsets, including CD4^+^ and CD8^+^ T cells (Figure 5B-C), NK cells (Figure 5D), CD11c^+^ MHCII^+^ dendritic cells (Figure 5E) and myeloid subsets expressing F4/80 (Figure 5F), Ly6G (Figure 5G), and Ly6C (Figure 5H). We similarly did not observe differences in numbers of CD45^lo^ microglia (Figure 5I), suggesting no major differences in microglial proliferation between genotypes in either region. These data suggested that, despite significant differences in the expression of major leukocyte chemoattractants in the cerebellum, differences in immune cell recruitment did not account for the increased pathogenesis observed in *Ripk3*^-/-^ mice during LGTV infection.

**Figure 5.**
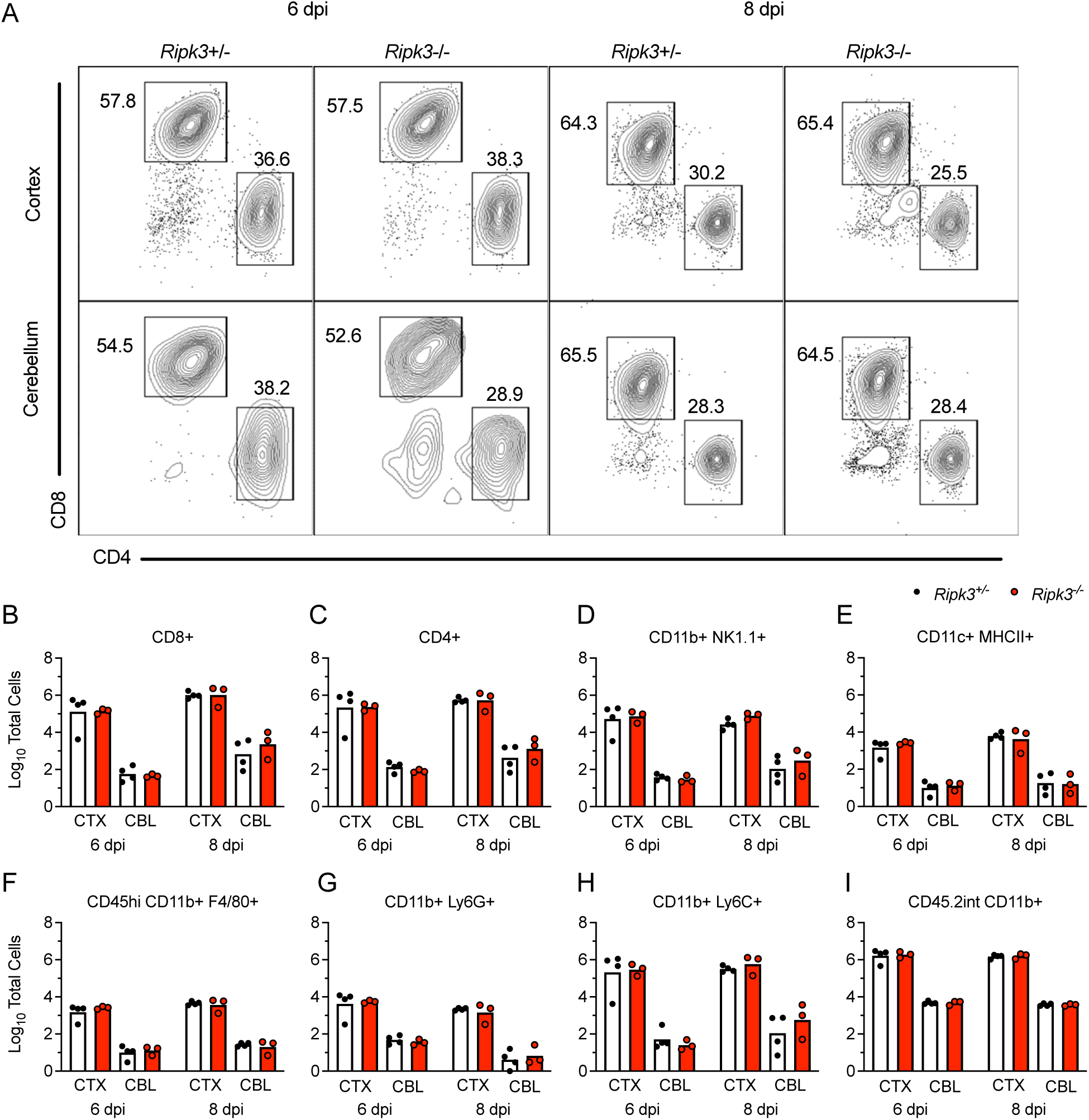
Leukocyte recruitment to the CNS occurs independently of RIPK3 signaling during LGTV encephalitis. (A-I) *Ripk3* ^-/-^ and littermate control mice were infected subcutaneously with LGTV TP21. Cerebral cortical and cerebellar tissues were harvested and leukocytes isolated for flow cytometric profiling at indicated days post infection (dpi). (A) Representative flow cytometry plots showing CD8+ and CD4+ T cells among CD3+ leukocytes in the brain. Numbers represent percentage of cells in each gate relative to total plotted cells. (B-I) Numbers of CD8+ T cells (B), CD4+ T cells (C), CD11b+ NK1.1+ natural killer cells (D), CD11c+ MHC-II+ dendritic cells (E), CD45^high^ CD11b+ F4/80+ macrophages (F), CD11b+ Ly6G+ neutrophils (G), CD45^high^ CD11b+ Ly6C+ monocytes (H), and CD45.2^lo^ CD11b+ microglia (I) among total brain leukocytes. No comparisons are statistically significant.

### RIPK3 promotes cell-intrinsic restriction of LGTV replication in cerebellar neurons

Given these observations, we next questioned whether *Ripk3*^-/-^ mice fail to control LGTV infection due to impaired innate immune restriction of LGTV replication. Assessment of viral burdens in brains of *Ripk3*^-/-^ mice following subcutaneous LGTV infection revealed that *Ripk3*^-/-^ mice exhibited significantly elevated CNS viral titers, particularly in the cerebellum, at both 8 and 12 dpi (Figure 6A). In contrast, *Mlkl*^-/-^ exhibited no such difference in viral burden in either brain region (Figure 6B). Differences in viral burden did not appear to be linked to deficits in blood-brain barrier integrity, as both *Ripk3*^-/-^ mice and littermate controls exhibited similar levels of sodium fluorescein extravasation into the CNS following infection (Figure 6C). We thus questioned whether RIPK3 was required for cell-intrinsic restriction of viral replication in susceptible CNS cell types. Multistep growth curve analysis in primary CNS cells revealed that pharmacologic inhibition of RIPK3 had no effect on LGTV replication in neurons derived from cerebral cortex (Figure 6D). In contrast, inhibition of RIPK3 significantly enhanced LGTV replication in primary cerebellar granule cell neurons cultures (Figure 6E). This effect was unique to neurons, as GSK 872 treatment had no impact on LGTV replication in primary astrocytes derived from either brain region (Figure 6F-G). Together, these data suggested that the enhanced pathogenesis observed in *Ripk3*^-/-^ mice was due to a specific failure to control infection in neurons of the cerebellum, resulting in enhanced overall CNS viral burden.

**Figure 6.**
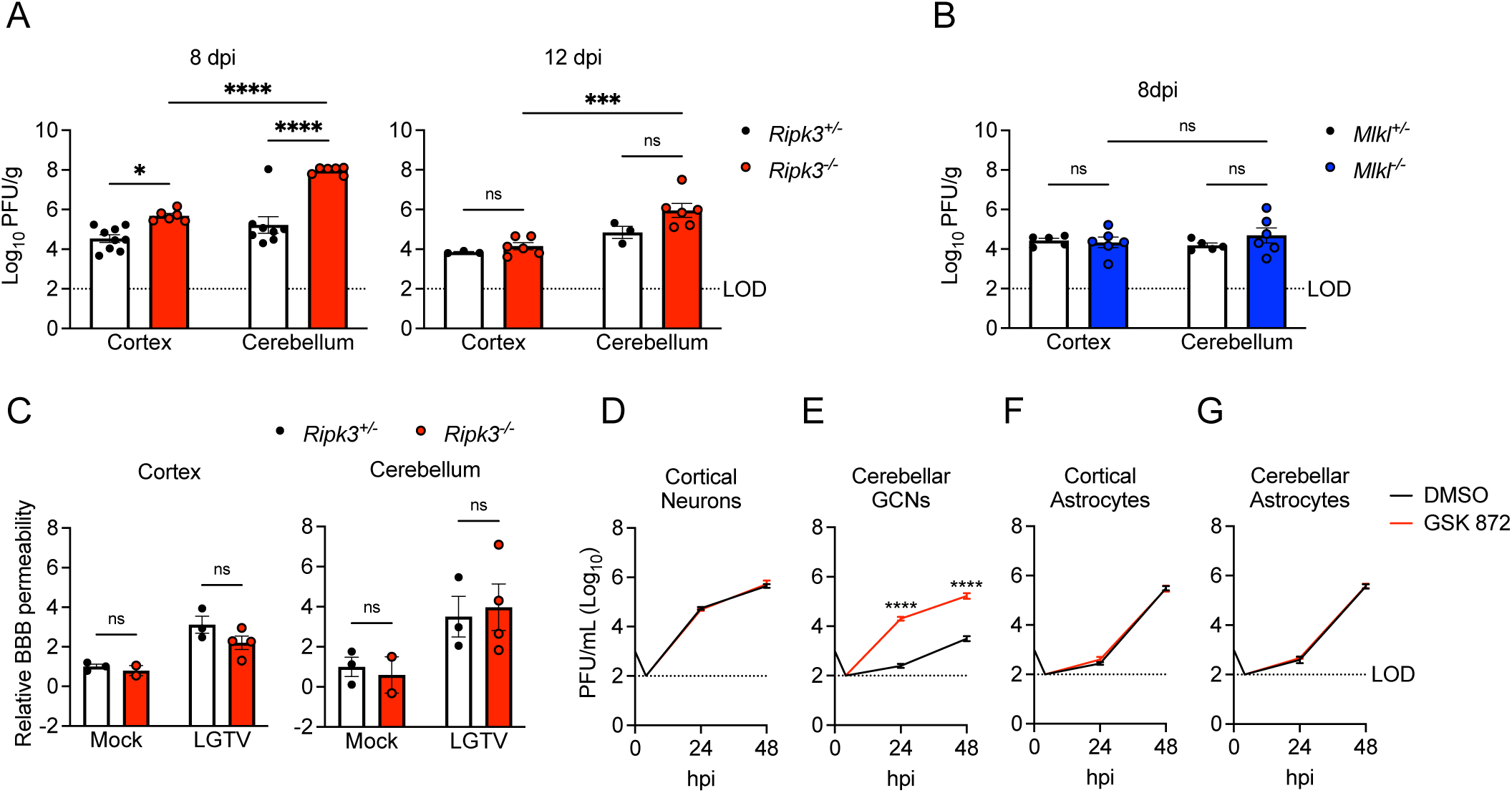
RIPK3 limits LGTV replication in cerebellar granule cell neurons. (A-B) *Ripk3* ^-/-^ (A) or *Mlkl* ^-/-^ (B) mice and littermate controls were infected subcutaneously with LGTV TP21. At 8 or 12 days post infection (dpi), viral loads in cerebral cortical and cerebellar tissues were determined by plaque assay. Data are pooled from 2-3 independent experiments. (C) *Ripk3* ^-/-^ and littermate control mice were subcutaneously infected with LGTV TP21. BBB permeability was measured at 8 dpi by detection of sodium fluorescein accumulation in tissue homogenates derived from cerebral cortex or cerebellum. Data represent individual brain fluorescence values normalized to serum sodium fluorescein concentration. Individual mouse values were then normalized to the mean values for uninfected controls. (D-G) Multistep growth curve analysis following infection with 0.01 MOI LGTV TP21 in cortical neurons (D), cerebellar granule cell neurons (E), cortical astrocytes (F), and cerebellar astrocytes (G). n=3 (cerebellar granule cell neurons) or 4 (astrocytes and cortical neurons) for growth curve experiments. ns, not significant. *p<0.05, **p < 0.01, ***p < 0.001, ****p < 0.0001. Error bars represent SEM.

### RIPK3 potentiates Type I IFN signaling in cerebellar neurons during LGTV infection

Our previous observation of diminished chemokine expression in cerebellar neurons derived from *Ripk3*^-/-^ mice suggested that these cells may exhibit broader deficits in innate immune signaling, resulting in poor control of LGTV replication. We, therefore, next questioned whether IFN signaling was perturbed in the cerebellum of mice lacking RIPK3 expression. Transcriptional profiling in brain tissues following subcutaneous LGTV infection revealed that, indeed, the cerebella of *Ripk3*^-/-^ mice exhibited diminished expression of many ISGs known to be critical for control of flavivirus replication (30–35), including *Ifit1, Isg15, Mx1, Mx2, Oas1b,* and *Rsad2,* while this phenotype was not observed in the cerebral cortex (Figure 7A-B). Similar analyses in primary cell cultures confirmed that cerebellar granule cell neurons, but not neurons derived from cerebral cortex, exhibited diminished expression of ISGs when RIPK3 signaling was blocked via GSK 872 treatment (Figure 7C-D). In contrast, we observed little to no impact of RIPK3 blockade on ISG expression in astrocytes derived from either brain region (Figure 7E-F). Together, these data demonstrate that RIPK3 signaling is required for the robust induction of type I IFN responses in neurons of the cerebellum, which is required for cell-intrinsic restriction of LGTV replication.

**Figure 7.**
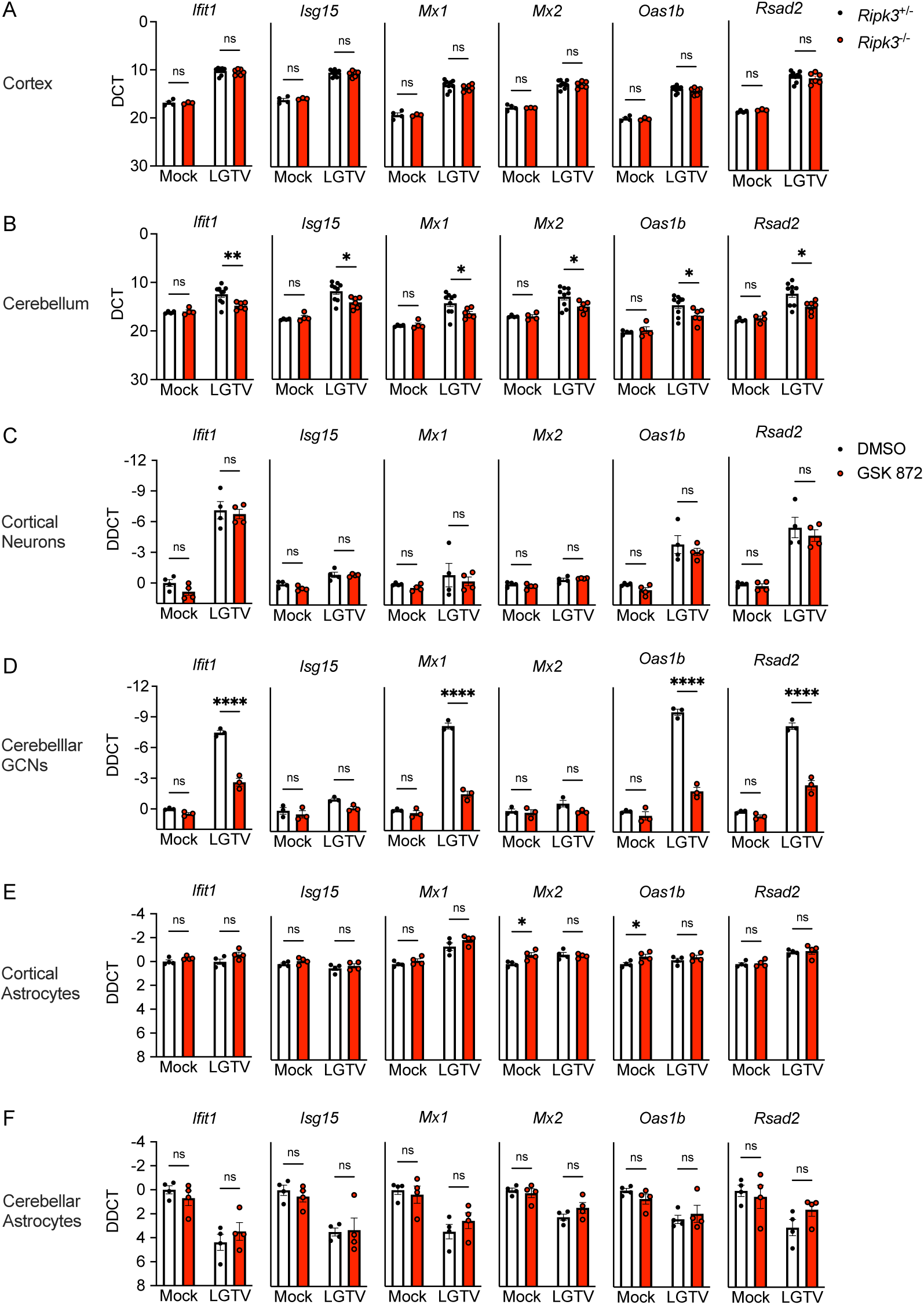
RIPK3 promotes ISG expression in cerebellar granule cell neurons. (A-B) *Ripk3* ^-/-^ and littermate control mice were infected subcutaneously with LGTV TP21. Transcriptional expression of indicated genes was assessed via qRT-PCR in cerebral cortical (A) and cerebellar (B) tissues at 8dpi. (C-D) Transcriptional expression of ISGs in wildtype (C57BL/6J) cultures of primary cortical neurons (C), cerebellar granule cell neurons (D), cortical astrocytes (E), and cerebellar astrocytes (F) following 2-hour pretreatment with GSK872 or vehicle and 24-hour infection with 0.5 (C-D) or 0.01 (E-F) MOI LGTV TP21, measured via qRT-PCR. ns, not significant. *p<0.05, **p < 0.01, ***p < 0.001, ****p < 0.0001. Error bars represent SEM.

To better understand the role of RIPK3 signaling in potentiating ISG expression, we next questioned whether RIPK3 acts downstream of IFN receptor signaling. To assess this, we treated neuron cultures with exogenous IFNβ for 1 hour following pretreatment with GSK 872 or vehicle control. As expected, IFNβ treatment resulted in robust induction of multiple ISGs (Figure 8A, Supplemental Figure 2A). However, pharmacologic blockade of RIPK3 did not impact ISG expression induced by IFNβ treatment in either cerebellar granule cell neurons (Figure 8A) or in cerebral cortical neurons (Supplemental Figure 2A), suggesting that RIPK3 likely does not act directly downstream of the type I IFN receptor (IFNAR) to modulate gene expression and/or that type I IFN alone is not sufficient to induce RIPK3 activation. We next tested the alternative hypothesis that RIPK3 regulates IFN signaling during LGTV infection by directly influencing the expression of IFN ligands and receptors. Surprisingly, transcriptional analysis revealed that pharmacologic blockade of RIPK3 did not influence the expression of the type I IFN ligands *Ifna* and *Ifnb* in cerebellar granule cell neurons following infection (Figure 8B). In contrast, GSK 843 treatment significantly blunted infection-induced upregulation of IFN receptor subunits, including the type I IFN receptor subunits *Ifnar1* and *Ifnar2*, and the type II IFN receptor subunits *Ifngr1*, and *Infgr2*. Other IFN ligands and receptors, including *Ifng* and the type III IFN ligands *Ifnl2* and *Ifnl3* were undetectable in all conditions (data not shown). Importantly, we did not observe this RIPK3-dependency in IFN receptor expression in cerebral cortical neuron cultures (Supplemental Figure 2B), suggesting that RIPK3 functions uniquely in cerebellar granule cell neurons to enhance type I IFN signaling during LGTV infection.

**Figure 8.**
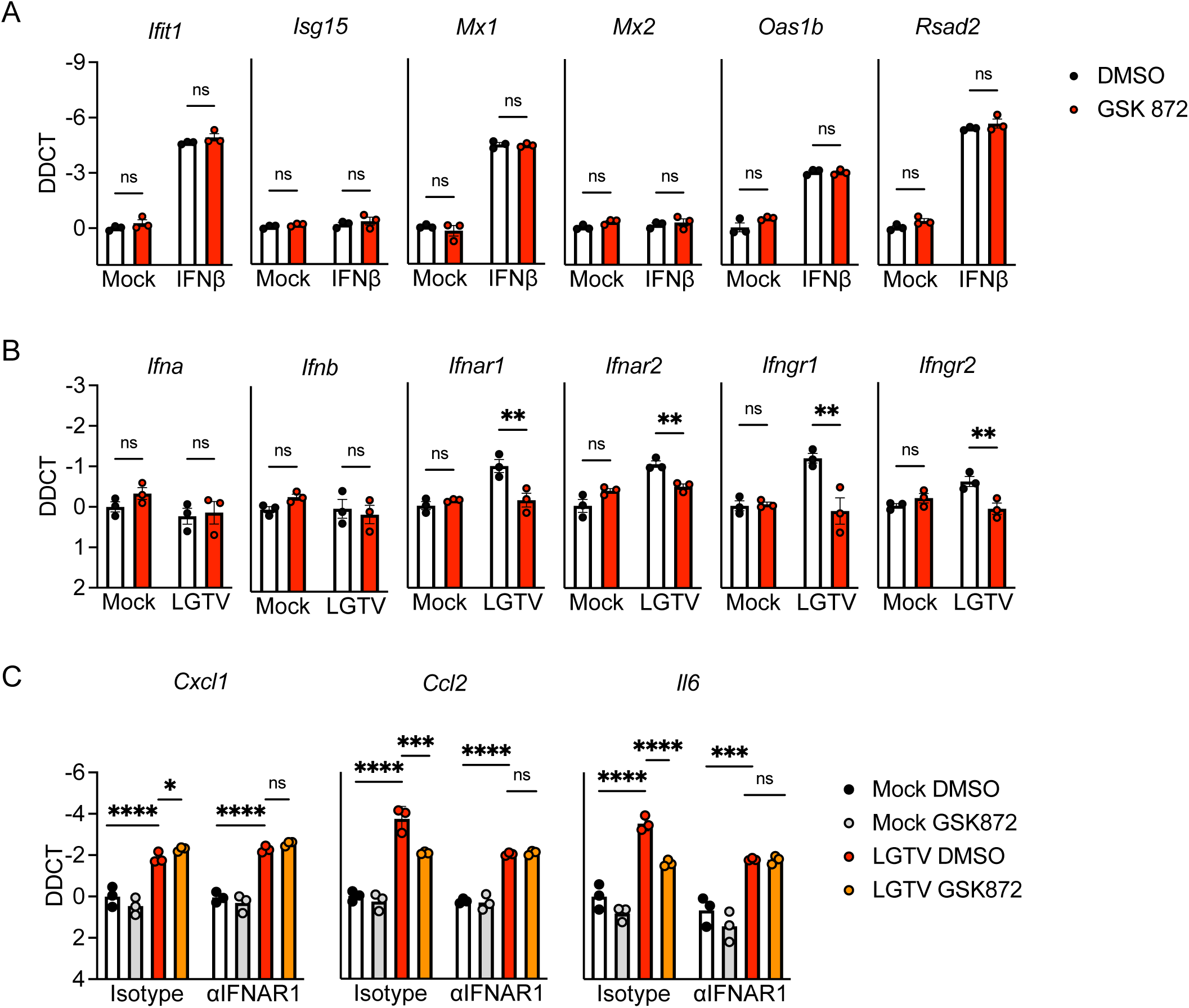
RIPK3 promotes expression of IFN receptors and IFN-dependent inflammatory genes in cerebellar granule cell neurons. A-B) Transcriptional expression of indicated genes in wildtype (C57BL/6J) cultures of cerebellar granule cell neurons in the setting of 2-hour pretreatment with GSK872 or vehicle followed by 1 hour treatment with 10ng/ml IFNβ (A) or 24-hour infection with 0.5 MOI LGTV TP21 (B). C) Expression of indicated genes in wildtype cerebellar granule cell neurons pretreated for 45 minutes with an anti-IFNAR1 neutralizing antibody or isotype control +/- cotreatment with GSK872 or vehicle, followed by 24-hour infection with 0.5 MOI LGTV TP21. ns, not significant. *p<0.05, **p < 0.01, ***p < 0.001, ****p < 0.0001. Error bars represent SEM.

To further investigate a role for RIPK3 in IFN-mediated gene expression in cerebellar neurons, we next performed experiments in which we blocked RIPK3 activity with or without simultaneous blockade of type I IFN signaling using a neutralizing antibody against IFNAR1. We reasoned that this paradigm would allow us to assess the differential influence of RIPK3 on IFNAR-dependent and IFNAR-independent gene expression following LGTV infection. Perhaps unsurprisingly, we observed that expression of most ISGs was completely dependent on IFNAR1 signaling, making it difficult to distinguish a specific role for RIPK3 in the absence of intact type I IFN signaling (Supplemental Figure 2C). We thus identified several alternative inflammatory genes whose expression was either completely (*Cxcl1*) or partially (*Ccl2 and Il6*) independent of IFNAR1 signaling following infection. Notably, pharmacologic blockade of RIPK3 only impacted the IFNAR1-depenent portion of the induced expression of these genes, while having no effect on the IFNAR1-independent portion, as indicated by a lack of effect in αIFNAR1-treated cultures (Figure 8C). Together, these data further support our observation of synergistic signaling between type I IFN and RIPK3 signaling in cerebellar granule cell neurons during LGTV infection.

## Discussion

Our findings identify a previously unknown function for RIPK3 in the coordination of brain region-specific innate immunity. The study of regional differences in neuroimmune signaling is a growing field, and there is accumulating evidence to suggest that resident neural cells exhibit differential responses to viral infection and cytokine stimulation across distinct anatomical regions of the CNS (36–39). Neurons and astrocytes in the cerebellum, in particular, have been shown to exhibit higher responsiveness to stimulation by type-I IFN, as well as to express higher basal levels of pathogen sensor molecules compared to other brain regions, suggesting a key evolutionary importance of innate antiviral defense in this tissue (12, 13). This regional difference in type I IFN signaling appears to underlie, at least in part, the relatively lower susceptibility of the cerebellum to flavivirus infection compared to susceptible regions of the forebrain, such as the cerebral cortex and hippocampus. However, the molecular mechanisms that determine the enhanced innate immune signaling observed in the cerebellum remain poorly understood. Our study suggests that RIPK3 signaling is required for the robust induction of ISG expression in cerebellar neurons during LGTV infection, although ongoing work is needed to understand the specific signaling interactions that mediate this effect.

Previous studies have described a highly complex interplay between RIPK3 and type I IFN signaling that varies significantly by cell type and disease model (17–19, 40). It is relatively clear that type I IFN signaling is capable of activating RIPK3 through various mechanisms, resulting in necroptosis and/or necroptosis-independent transcriptional activation (40–43). However, how RIPK3 operates *upstream* of (or synergistically with) type I IFN signaling to influence expression of ISGs is less clear. We and others have shown that ISG expression is significantly diminished in a variety of settings when RIPK3 signaling is ablated (17, 18), including in cerebellar granule cell neurons during LGTV infection in this study. One possible explanation for this effect is RIPK3-mediated activation of NF-κB, a transcription factor strongly associated with RIPK signaling with known roles in potentiating type I IFN signaling and ISG expression (22, 44–46). We and others also previously showed that RIPK3 activation in cortical neurons following ZIKV infection leads to interferon regulatory factor 1 (IRF1) activation, which was required for expression of at least a subset of RIPK3-induced genes in that setting, although this effect is likely indirect, as IRF1 is not a known RIPK3 substrate (17). Additional work will be needed to fully characterize the regulatory mechanisms that are invoked in the interplay between RIPK3 and type I IFN signaling in the CNS.

Our study also further expands our understanding of the necroptosis-independent functions for RIPK3 signaling in the CNS. Many studies have now firmly established the importance of RIPK3 in promoting host defense through mechanisms independent of its canonical role in necroptosis (16–21). However, these necroptosis-independent functions appear to vary significantly by disease state, including CNS infection with distinct neuroinvasive flaviviruses (47, 48). We and others previously showed that the primary role for RIPK3 in restricting WNV encephalitis was the induction of chemokine expression and the recruitment of antiviral leukocytes into the infected CNS (16). Notably, while we did observe RIPK3-mediated chemokine expression in the cerebellum during LGTV infection, this chemokine expression was apparently dispensable for CNS immune cell recruitment. Instead, the transcriptional activation of antiviral effector genes, including ISGs, was required for cell-intrinsic restriction of LGTV replication in neurons, a phenotype more similar to our findings with ZIKV (17), although we did not observe evidence for a regional specification of this response during ZIKV infection. In contrast to these observations, Bian and colleagues have observed quite distinct phenotypes in a model of JEV encephalitis, wherein both RIPK3 and MLKL appeared to exacerbate rather than restrict disease pathogenesis (49, 50). RIPK3 also appeared to *suppress* rather than promote ISG expression in JEV infected neurons. The factors that determine such distinct outcomes of RIPK3 signaling across this family of closely related viruses are mysterious and are the subject of ongoing investigation by our laboratory and others.

## Materials and Methods

### Mouse lines

*Ripk3*^-/-^ (51) *Mlkl*^-/-^ (52) mouse lines were bred and housed under specific-pathogen free conditions in Nelson Biological Laboratories at Rutgers University. *Ripk3*^-/-^ mice were generously provided by Genentech, Inc. Wild-type C57BL/6J mice were either obtained commercially (Jackson Laboratories) or bred in-house. Mice used for subcutaneous infections were 5 weeks old; mice used for intracranial infections were 8-15 weeks old.

### Virus and titer determination

Langat virus strain TP21 was used throughout the study. Founder stocks were obtained from the World Reference Center for Emerging Viruses and Arboviruses (WRCEVA). Laboratory stocks were generated using Vero E6 cells (ATCC, #CRL-1586) and frozen at −80°C until needed. Virus titers were determined by plaque assay on Vero E6 cells. Cells were maintained in DMEM (Corning #10-013-CV) supplemented with 10% Heat Inactivated FBS (Gemini Biosciences #100-106), 1% Penicillin–Streptomycin-Glutamine (Gemini Biosciences #400-110), 1% Amphotericin B (Gemini Biosciences #400–104), 1% Non-Essential Amino Acids (Cytiva, #SH30238.01), and 1% HEPES (Cytiva, #SH30237.01). Plaque assay media was composed of 1X EMEM (Lonza # 12-684F) supplemented with 2% Heat Inactivated FBS (Gemini Biosciences #100-106), 1% Penicillin–Streptomycin-Glutamine (Gemini Biosciences, #400-110), 1% Amphotericin B (Gemini Biosciences #400-104), 1% Non-Essential Amino Acids (Cytiva, #SH30238.01), and 1% HEPES (Cytiva, SH30237.01), 0.75% Sodium Bicarbonate (VWR, #BDH9280) and 0.5% Methyl Cellulose (VWR, #K390). Plaque assays were developed at 5dpi by removal of overlay media and staining/fixation using 10% neutral buffered formalin (VWR, #89370) and 0.25% crystal violet (VWR, #0528). Plaque assays were performed by adding 100uL of serially diluted sample for 1 hour at 37°C to 12-well plates containing 200,000 Vero E6 cells per well. Plates were further incubated with plaque assay media at 37°C and 5% CO2 for 5 days. Medium was removed from the wells and replaced with fixative containing crystal violet for approximately 20-30 minutes. Plates were washed repeatedly in H_2_O and allowed to dry before counting visible plaques.

### Mouse infections and tissue harvesting

Isoflurane anesthesia was used for all procedures. Mice were inoculated subcutaneously (50uL) with 3×10^4^ PFU or injected intracranially (10uL) with 50 PFU of LGTV-TP21 using insulin syringes (BD Medical, #BD-329461). At appropriate times post infection, mice underwent cardiac perfusions with 30 mL cold sterile 1X phosphate-buffered saline (PBS). Extracted tissues were weighed and homogenized using 1.0 mm diameter zirconia/silica beads (Biospec Products, #11079110z) in sterile PBS for plaque assay or TRI Reagent (Zymo, #R2050-1) for gene expression analysis. Homogenization was performed in an Omni Beadrupter Elite for 2 sequential cycles of 20 s at a speed of 4 m/s.

### Primary cell infections

Cortical and cerebellar astrocytes were harvested from P1-P2 pups and cortical neurons were harvested at E13.5-E15.5. Tissues were dissociated using the Neural Dissociation Kit (T) following manufacturer’s instructions (Miltenyi, #130-093-231). Astrocytes were expanded in AM-a medium (ScienCell, #1831) supplemented with 10% FBS in fibronectin-coated cell culture flasks and seeded into plates coated with 20 μg/mL Poly-L-Lysine (Sigma-Aldrich, #9155) before experiments. Neurons were seeded into PLL-coated cell culture treated plates and grown in Neurobasal Plus + B-27 supplement medium (Thermo-Fisher Scientific, #A3582901) prior to use in experiments 7-9 days in vitro (DIV). Mouse cerebellar granule cells from C57BL/6 mice (ScienCell, # M1530-57) were seeded into cell culture treated plates coated with 10 ug/mL Poly-D-Lysine (ThermoFisher, #A3890401) containing prewarmed Neuronal Medium (ScienCell, #1521) following manufacturer recommendations and used for experiments 6 DIV.

Macrophages and dendritic cells were isolated from bone marrow of euthanized mice. Femurs were isolated and bone marrow pushed out using a sterile needle and syringe loaded with RPMI supplemented with 10% FBS, 1% Penicillin–Streptomycin-Glutamine, 1% HEPES, 1% Glutamax (ThermoFisher, #35050061). Bone marrow was plated into non-cell-culture treated 10cm petri dishes in 8mL supplemented RPMI medium containing either 20ng/mL recombinant M-CSF (Peprotech, #315-02) or 20ng/mL recombinant GM-CSF (Peprotech, #315-03) and 20ng/mL IL-4 (Peprotech, #214-14) for differentiation into macrophages or dendritic cells, respectively. Cells were fed with additional medium containing the appropriate cytokines four days later and used for experiments at 6-7 DIV. Cells were seeded into cell-culture treated dishes prior to experimentation. For viral replication determination, all cultures were infected with LGTV TP21 at an MOI of 0.01. For qRT-PCR experiments, cortical and cerebellar neuron cultures were infected at an MOI of 0.5, while astrocyte cultures were infected using an MOI of 0.01. The pharmacologic inhibitor of RIPK3, GSK872, was added to cultures at 100nM for 2 hours prior to infection or subsequent treatments. Interferon-β was added to neuron cultures at 10ng/mL for 1 hour prior to harvesting cell lysates. IFNAR-1 monoclonal antibody (MAR1-5A3, Leinco Technologies) or isotype control (GIR-208, Leinco Technologies) were added to cultures at 5µg/mL 45 minutes prior to Langat virus infection.

### Quantitative real-time PCR

Total RNA from harvested tissues was extracted using Zymo Direct-zol RNA Miniprep kit, as per manufacturer instructions (Zymo, #R2051). Total RNA extraction from cultured cells, cDNA synthesis, and subsequent qRT-PCR were performed as previously described (22, 53). Cycle threshold (CT) values for analyzed genes were normalized to CT values of the housekeeping gene 18 S (CT_Target_ − CT_18S_ = ΔCT). Data from primary cell culture experiments were further normalized to baseline control values (ΔCT_experimental_ − ΔCT_control_ = ΔΔCT (DDCT)). A list of primers used in this study can be found in **Supplemental Table 1**.

### Flow Cytometry

The cerebella and cerebral cortices of mouse brains were dissected from freshly perfused mice and placed into tubes containing 1X PBS. Brain tissues were incubated with 10mL buffer containing 0.05% Collagenase Type I (Sigma-Aldrich, #C0130), 10ug/mL DNase I (Sigma-Aldrich, #D4527) and 10mM HEPES (Cytiva, #SH30237.01) in 1X Hanks’ Balanced Salt Solution (VWR, #02-1231-0500) for one hour at room temperature under constant rotation. Brain tissues were transferred to a 70um strainer on 50mL conical tubes and mashed through the strainer using the plunger of 3-5mL syringes. Tissue was separated in 8 mL 37% Isotonic Percoll (Percoll: Cytiva, #17-0891-02; RPMI 1640: Corning, #10-040-CV, supplemented with 5% FBS) by centrifugation at 1200xg for 30 minutes with a slow break. The myelin layer and supernatant were discarded. Leukocytes were incubated in 1X RBC Lysis Buffer (Tonbo Biosciences, #TNB-4300-L100) for 10 minutes at room temperature. Cells were centrifuged and resuspended in FACS buffer composed of 1X PBS, 2% sodium azide and 5% FBS. Samples were transferred into a U-bottomed 96-well plate. Leukocytes were blocked with 2% normal mouse serum and 1% FcX Block (BioLegend, #101320) in FACS buffer for 30 minutes at 4°C prior to being stained with fluorescently conjugated antibodies to CD3e (Biolegend, clone 17A2), CD44 (Biolegend, clone IM7), CD19 (Biolegend, clone 6D5), CD8a (Biolegend, clone 53-6.7), CD4 (Biolegend, clone RM4-5), CD45.2 (Biolegend, clone 104), MHC-II (Biolegend, clone M5/114.15.2), NK1.1 (Biolegend, clone PK136), CD11c (Biolegend, clone N418), F4/80 (Biolegend, clone BM8), CD11b (Biolegend, clone M1/70), Ly6G (Biolegend, clone 1A8), Ly6C (Biolegend, clone HK1.4), CD80 (Biolegend, clone 16-10A1), and Zombie NIR (Biolegend, #423105). Leukocytes were stained for 30 minutes at 4C prior to washing in FACS buffer and fixation with 1% PFA in PBS (ThermoFisher, #J19943-K2). Data collection and analysis were performed using a Cytek Northern Lights Cytometer (Cytek, Fremont, California) and FlowJo software (Treestar). Data were normalized using a standard bead concentration counted by the cytometer with each sample (ThermoFisher, #C36950). Spleens were crushed between two slides, filtered through a 70um cell strainer, and washed with FACS buffer. Isolated splenocytes were incubated with 1X RBC Lysis Buffer as done for leukocytes isolated from the brain prior to blocking and staining.

### In vivo assessment of blood brain barrier permeability

In vivo assessment of blood brain barrier permeability was carried out as described (54).,Mice were injected intraperitoneally with 100uL of 100mg/mL fluorescein sodium salt (Sigma, #F6377) dissolved in sterile 1X PBS. After 45 minutes, blood was collected followed by cardiac perfusion. Tissues were dissected and homogenized in 1X PBS as described above. Serum and supernatant from homogenized tissues were incubated overnight at 4°C with 2% Trichloroacetic acid solution (Sigma, #T0699) at a 1:1 dilution. Precipitated protein was pelleted by 10 minutes of centrifugation at 2,823xg at 4°C. Supernatants were diluted with borate buffer, pH 11 (Sigma, #1094621000) to achieve a neutral pH. Fluorescein emission at 538nm was measured for samples in an optically clear black-walled 96-well plate (Corning, #3904) using a SpectraMax iD3 plate reader (Molecular Devices, San Jose, CA). Tissue fluorescence values were standardized against plasma values for individual mice.

### Statistical analysis

Normally distributed data were analyzed using appropriate parametric tests: two-way analysis of variance (ANOVA) with Sidak’s correction for multiple comparisons and Log-rank (Mantel-Cox) test for survival comparison, both using GraphPad Prism Software v8 (GraphPad Software, San Diego, CA). Chi square tests for comparison of clinical disease signs was performed using Excel v2211 (Microsoft). P < 0.05 was considered statistically significant.

## Acknowledgements

This work was supported by R01 NS120895 (to BPD). JPA and IE were supported by NIH Supplement to Promote Diversity (R01 NS120895-S1 and NS120895-S2). NPC was supported by F31 NS124242.

---

**Supplemental Figure 1:**
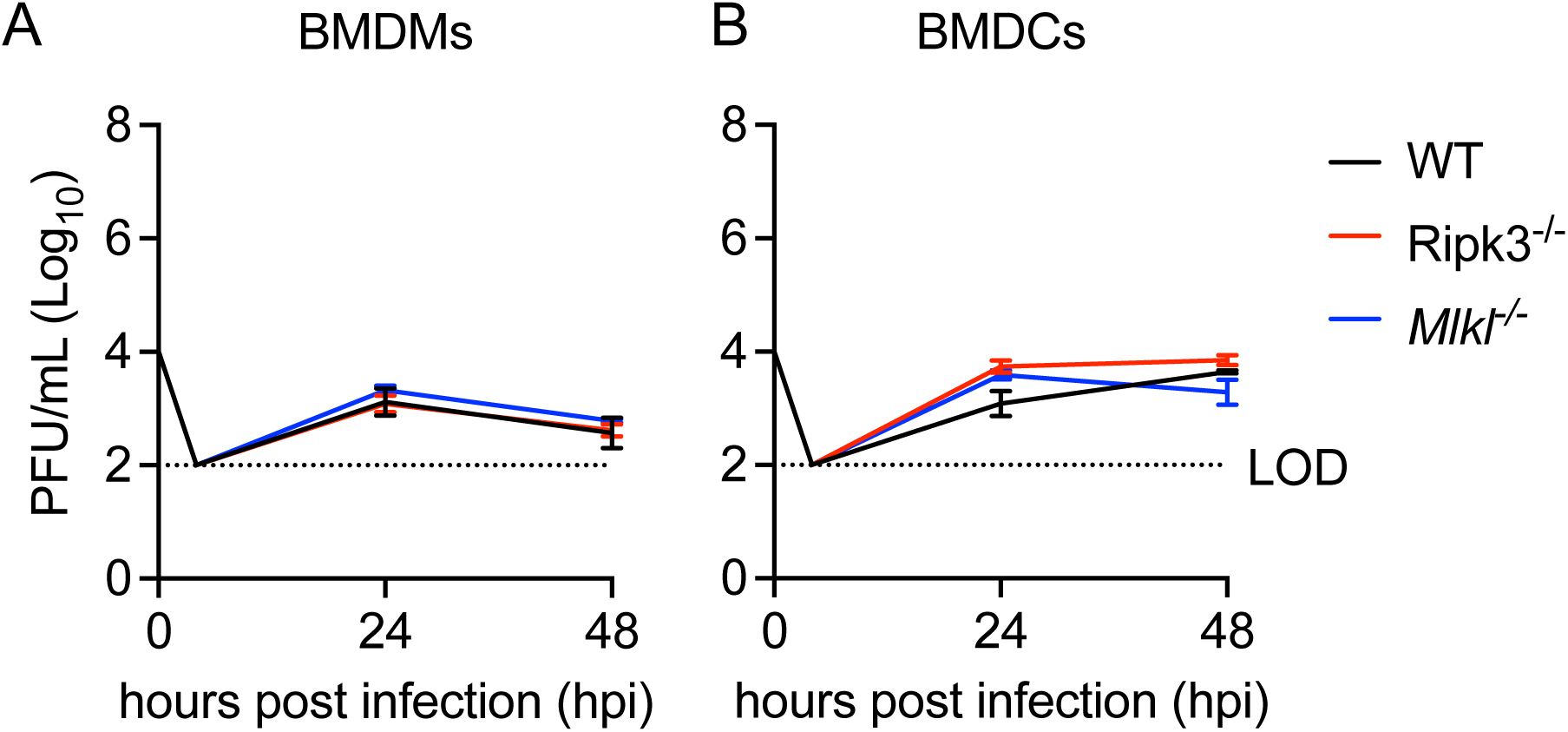
Neither RIPK3 nor MLKL is required for restriction of LGTV replication in bone marrow-derived macrophages and dendritic cells. (A-B) Multistep growth curve analysis following infection with 0.01 MOI LGTV TP21 in primary macrophages (BMDMs) (A) and dendritic cells (BMDCs) (B) cultured from bone marrow of C57BL/6J (WT), *Ripk3*^-/-^, or *Mlkl*^-/-^ mice. (n=4) No comparisons are statistically significant.

**Supplemental Figure 2.**
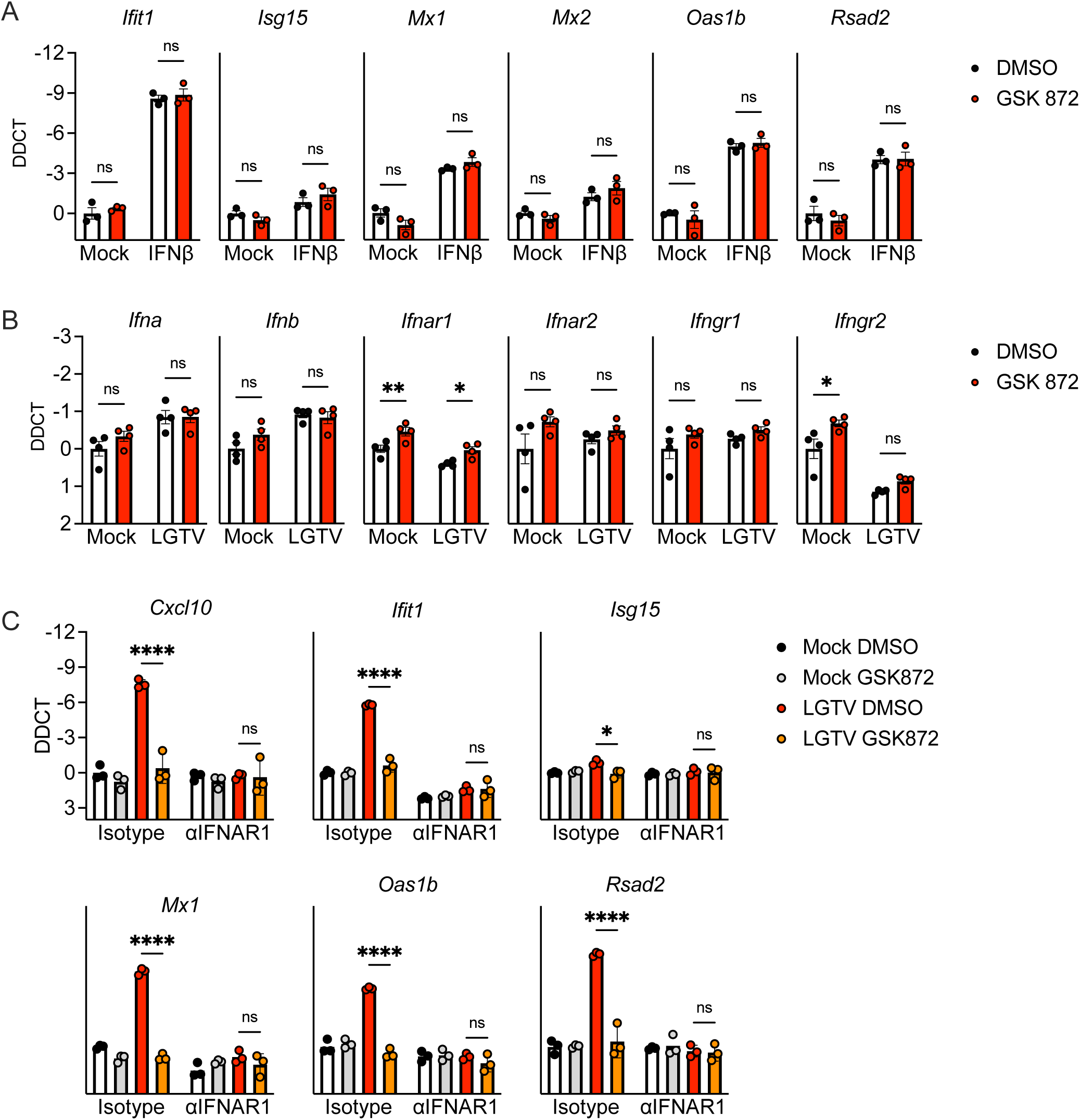
**RIPK3 does not impact IFN-mediated responses to LGTV in cortical neurons. A-B**) Transcriptional expression of indicated genes in wildtype (C57BL/6J) cultures of cerebral cortical neurons in the setting of 2-hour pretreatment with GSK872 or vehicle followed by 1 hour treatment with 10ng/ml IFNβ (A) or 24-hour infection with 0.5 MOI LGTV TP21 (B). C) Expression of indicated genes in wildtype cerebral cortical neurons pretreated for 45 minutes with an anti-IFNAR1 neutralizing antibody or isotype control +/- cotreatment with GSK872 or vehicle, followed by 24-hour infection with 0.5 MOI LGTV TP21. ns, not significant. *p<0.05, **p < 0.01, ***p < 0.001, ****p < 0.0001. Error bars represent SEM.

**Supplemental Table 1:**
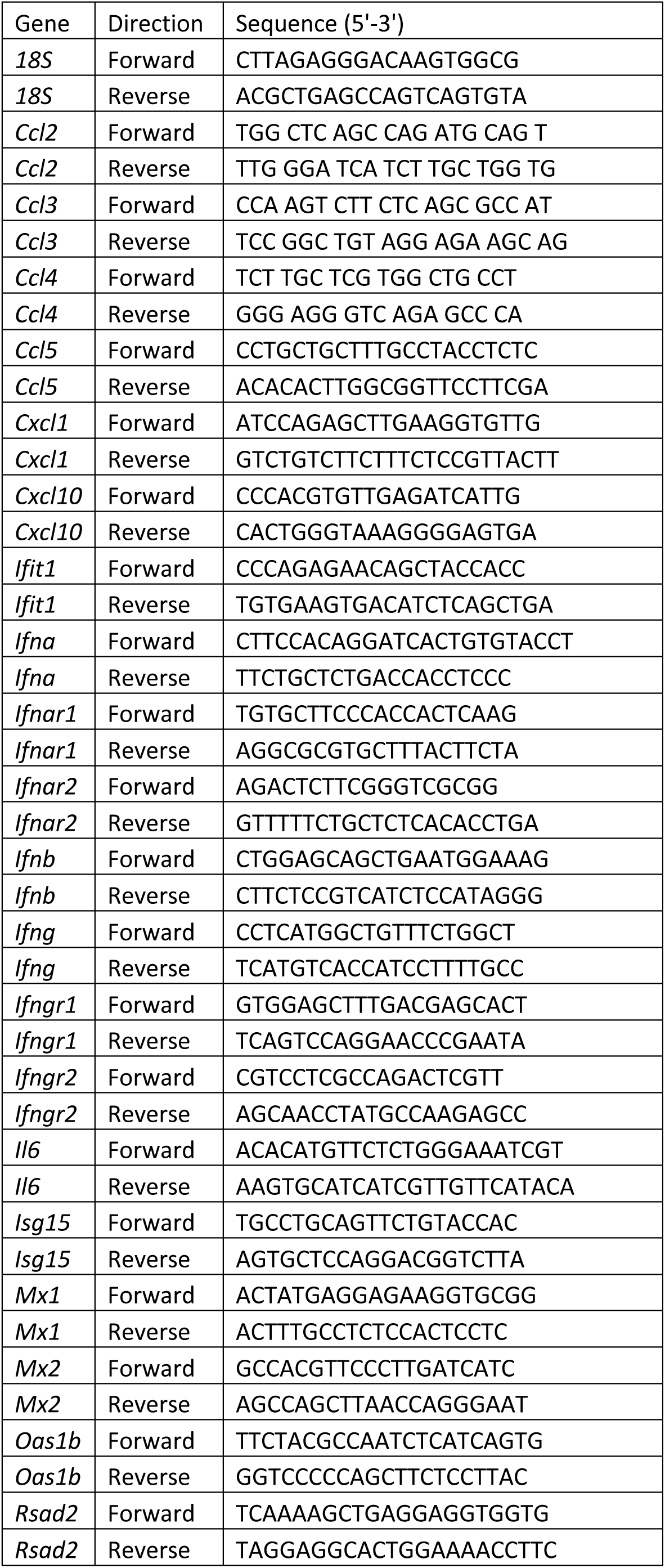
Primer sequences for qRT-PCR.

